# Resolving the source of branch length variation in the Y chromosome phylogeny

**DOI:** 10.1101/2024.07.05.602100

**Authors:** Yaniv Swiel, Janet Kelso, Stéphane Peyrégne

**Affiliations:** Department of Evolutionary Genetics, Max Planck Institute for Evolutionary Anthropology, Leipzig, Germany

**Keywords:** Reference bias, Ancient DNA, Y chromosome, Sequence alignment, Molecular dating, Mutation rate, Generation time

## Abstract

Genetic variation in the non-recombining part of the human Y chromosome has provided important insight into the paternal history of human populations. However, a significant and yet unexplained branch length variation of Y chromosome lineages has been observed, notably amongst those that are highly diverged from the human reference Y chromosome. Understanding the origin of this variation, which has previously been attributed to changes in generation time, mutation rate, or efficacy of selection, is important for accurately reconstructing human evolutionary and demographic history.

Here, we analyze Y chromosomes from present-day and ancient modern humans, as well as Neandertals, and show that branch length variation amongst human Y chromosomes cannot solely be explained by differences in demographic or biological processes. Instead, reference bias results in mutations being missed on Y chromosomes that are highly diverged from the reference used for alignment. We show that masking fast-evolving, highly divergent regions of the human Y chromosome mitigates the effect of this bias and enables more accurate determination of branch lengths in the Y chromosome phylogeny. Finally, we show that this approach allows us to estimate the age of ancient samples from Y chromosome sequence data and provide updated TMRCA estimates using the portion of the Y chromosome where the effect of reference bias is minimised.

## Introduction

Human Y chromosome lineages that are highly diverged from the reference Y chromosome appear to have accumulated fewer mutations than lineages that are more closely related to the reference, a non-African lineage of the R1b haplogroup. In particular, some human Y lineages in Africa, such as A00 [1], appear to have accumulated fewer mutations than other Y lineages since they last shared a common ancestor. In their phylogenetic analysis of a diverse selection of human Y chromosomes for worldwide populations, Wei *et al.* [2] observed a shortened branch for the A3 haplogroup represented by a single individual in their study. They attributed this shortened branch to an under-calling of variants due to the low-coverage sequencing (5x) of the A3 individual. Scozzari *et al.* [3] analysed Y chromosomes representing major haplogroups and noted shortened branches for the A haplogroups relative to the rest of the tree. They were not able to determine the cause of the shortened branches, but suggested that a reduction in the male effective population size coinciding with the out-of-Africa bottleneck allowed mildly deleterious mutations to accumulate on non-African Y chromosomes. Hallast *et al.* [4] tested whether the tissue from which the DNA is extracted has an effect on the branch length heterogeneity but could not detect any statistically significant differences between tissues. They therefore proposed that a change in the mean paternal age (resulting in the increase or decrease of the effective mutation rate) in a particular geographical region could result in associated haplogroups displaying similarly shortened or lengthened branches. Barbieri *et al.* [5] suggested a number of technical biases that could result in the undercalling of variants in A and B Y chromosome lineages. They proposed that capture bias could be introduced when using a Y chromosome capture array based on the reference Y chromosome and that the underrepresentation of A and B haplogroups in reference datasets could affect genotype calling and/or imputation. They also suggested that, in addition to these biases, population-based differences in average paternal age could contribute to the branch length variation across the Y chromosome phylogeny through the modification of the mutation rate. Although these studies relied on capture sequencing data (and low-coverage shotgun sequencing data in the case of Wei *et al.* [2]), Naidoo *et al.* [6] also noticed shortened branches among A haplogroups using high-coverage shotgun sequencing data, suggesting that these differences do not originate from capture bias.

Understanding the cause of branch length variation in the Y chromosome phylogeny is important to accurately reconstruct human evolutionary history. The A00 Y chromosome lineage has been used to estimate the time to the most recent common ancestor (TMRCA) of the Y chromosomes of all living humans [7] and studies of archaic human Y chromosomes [1, 8] have used this TMRCA to estimate the TMRCA between modern and archaic human Y chromosomes. The branch length variation observed suggests that there may be differences in mutation rate among Y chromosome lineages. Therefore, incorrectly assuming a constant mutation rate over the Y chromosome phylogeny could result in biased TMRCA estimates that do not accurately reflect population split times. Moreover, variation in the Y chromosome mutation rate could bias methods that estimate the age of ancient and archaic humans by comparing the number of mutations accumulated on their Y chromosomes to those of present-day human Y chromosomes.

Newly sequenced, high-quality Y chromosomes from present-day and ancient modern humans, as well as from Neandertals, provide an opportunity to re-evaluate the hypotheses that changes in mutation rate or generation time are the cause of the branch length variation in the Y chromosome phylogeny. These Y chromosomes also allow us to test the effect of reference bias on the mapping of short reads. Reference bias refers to the observation that sequencing reads carrying non-reference alleles are less likely to align successfully to the reference genome than those carrying reference alleles. The effect of reference bias is exacerbated for shorter read lengths and for reads carrying ancient DNA damage, which introduces more mismatches to the reference genome [9, 10]. Y chromosomes that are divergent from the reference Y chromosome may be particularly affected by reference bias, making it a possible explanation for the branch shortening reported for some African Y chromosome lineages. The recent availability of high-quality Y chromosome assemblies — including a telomere-to-telomere Y chromosome assembly [11] and an almost fully contiguous assembly of a Y chromosome representing the basal A0b haplogroup [12] — allow us to test whether the reference used for alignment has any effect on branch length.

## Results

### Branch length variation among a diverse set of Y chromosomes

To understand why there appear to be fewer mutations in Y chromosomes that are diverged from the reference used for alignment, we compiled a dataset comprising high-quality Y chromosomes from both present-day and ancient modern humans, as well as from Neandertals (Table S1). While the dataset contains Y chromosomes representing all of the major haplogroups, our focus was on incorporating Y chromosomes that are significantly diverged from the human reference (hg19) Y chromosome. To determine whether there is branch length variation in our dataset, we compared the number of apparent mutations on each Y chromosome to the number in a present-day non-African Y chromosome (haplogroup O1b1a1a1a) since their common ancestor, thereafter referred to as relative branch length difference. If the mutation rate was constant across the Y chromosome phylogeny, we would expect no significant differences between the number of mutations that we detect on the Y chromosomes of present-day individuals. However, significantly fewer mutations are identified for those African Y chromosomes that are the most diverged from the reference, and this difference increases linearly with increasing divergence from the reference (Figure 1). We find that *∼*55 mutations are missed with each additional pairwise difference from the reference per 10,000 basepairs (bp) (Figure S1).

**Fig. 1:**
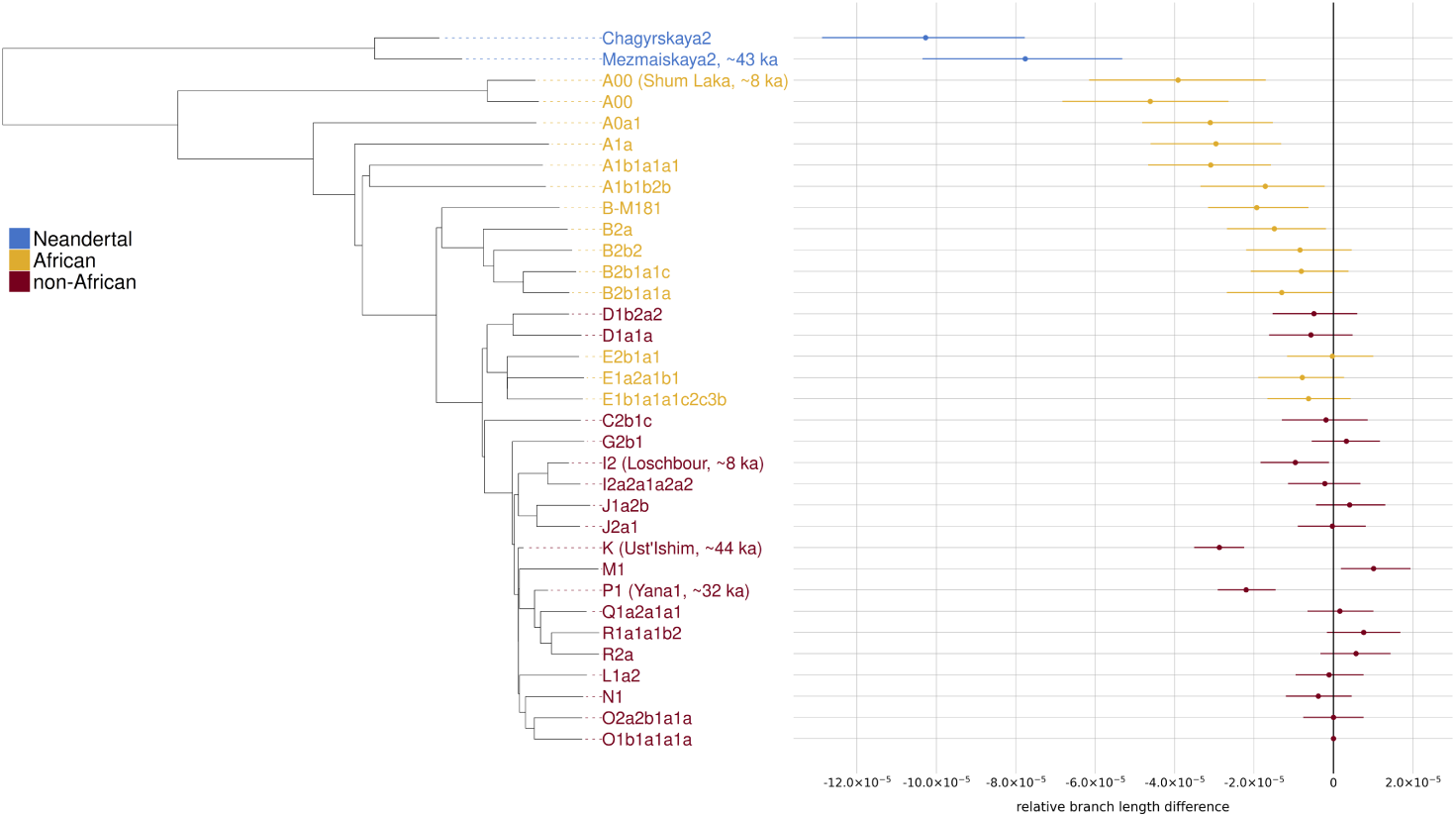
Neighbour-joining tree of a Y chromosome phylogeny comprising two Neandertals [1, 13], four ancient humans [14–17] and twenty eight present-day humans [18, 19] with associated relative branch length differences (i.e. the difference in branch length normalised by the total number of sites used for the comparison) compared to a present-day non-African Y chromosome (O1b1a1a1a). The colours indicate the population of origin. The error bars represent 95% confidence intervals (CIs) computed by resampling branch lengths from a Poisson distribution as described in Petr *et al.* [1].

We expect to observe shorter branches for ancient individuals than for modern individuals because they have less time to accumulate mutations. This explains the shorter branches for *Ust’Ishim* and *Yana1* (Figure 1). However, we would expect that ancient individuals with similar radiocarbon dates should have branch lengths similar to one another. Instead, when we compare the Y chromosomes of ancient individuals with similar radiocarbon dates but with differing levels of divergence from the reference (i.e. *Mezmaiskaya2* vs *Ust’Ishim*/*Shum Laka* vs *Loschbour*), we find that the divergent branches are significantly shorter than the non-divergent branches. As shortened branches are specific to some African haplogroups (i.e. those most diverged from the reference) rather than to all Africans (Figure 1), this branch length variation is unlikely to be the result of an accumulation of mutations on non-African Y chromosomes following the out-of-Africa bottleneck as proposed by Scozzari *et al.* [3].

### Testing the hypothesis of mutation rate and generation time variation

The significant branch length variation observed over the Y chromosome phylogeny suggests that the assumption of a constant Y chromosome mutation rate and constant generation time may be violated. Therefore, we next estimated the changes in mutation rate and generation time required to explain the observed differences in branch lengths. Previous studies [1, 14] have estimated the Y chromosome mutation rate based on the difference in the number of mutations between an ancient human Y chromosome (that of a 44.5 thousand year old Eurasian hunter-gatherer, *Ust’Ishim*) [14] and those of present-day humans. Starting with the mutation rate of 7.34*×*10*^−^*^10^ mutations/bp/year (which was computed from present-day non-African Y chromosomes [1]), we conducted a series of pairwise branch length comparisons to estimate the relative changes in mutation rate that would result in the observed branch length differences. We used the Y chromosomes that are the most diverged from the reference and calibrated our estimates according to the known ages of the individuals. We find that multiple changes in mutation rate are needed to explain the data and that, in total, the mutation rate would need to have increased by 66% since the split between humans and Neandertals (Figure 2a and Table S2). This contrasts with the finding, based on a comparison between the complete assemblies of human and chimpanzee Y chromosomes, that there has been an 11% decrease in the human Y chromosome mutation rate since the divergence between humans and chimpanzees *∼*6 million years ago [20]. Mutation rate differences alone are, therefore, unlikely to explain the observations.

**Fig. 2:**
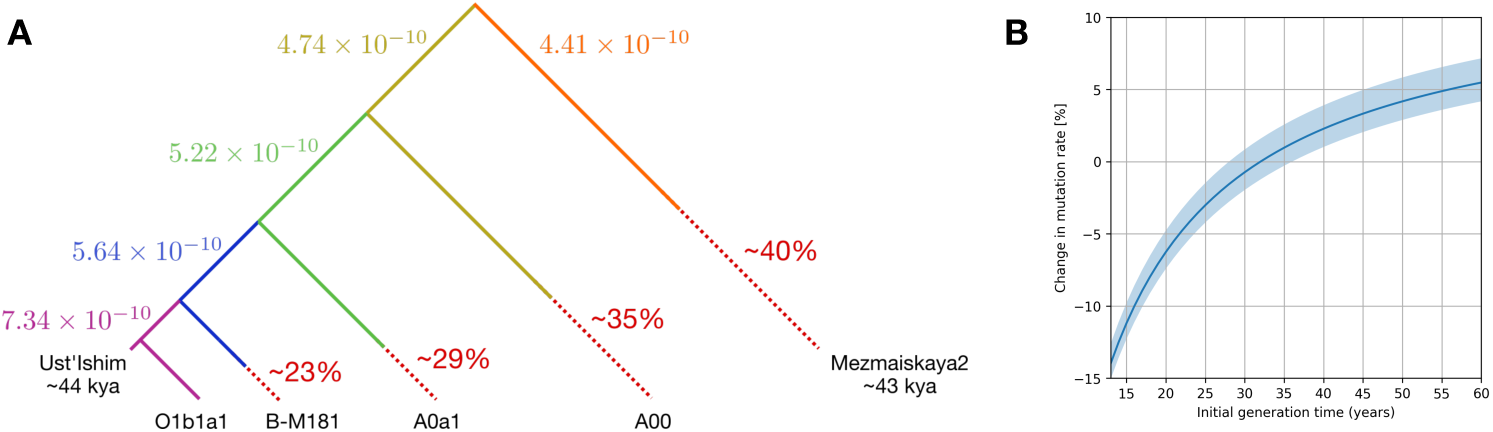
(a) Mutation rates (in units of mutations/bp/year) required to explain the branch lengths of the Y chromosomes that are highly diverged from the human reference. The dashed lines represent the branch shortening relative to the O1b1a1a1a branch. The colours indicate the branches of the tree and their corresponding mutation rate. (b) Percentage change in the mutation rate if the male generation time changed from the value on the x-axis to 32 years (the current average estimate in present-day populations [22]). The shaded area corresponds to the change in mutation rate if the current generation time is instead between 28 and 36 years.

While the *de novo* mutation rate increases with paternal age (see Methods), a decrease in generation time (i.e. in the average paternal age) can result in an increase in the number of mutations because more generations occur over time. To determine if the branch length variation could be caused by changes in male generation time, we estimated what change in generation time could have lead to a 66% increase in mutation rate (Figure 2b, see Methods). We relied on the relationship between paternal age and autosomal mutation rate as this relationship for the Y chromosome mutation rate is difficult to accurately estimate. This is due to the small size of the Y chromosome, which accumulates relatively few mutations per generation, and the limitations of short read sequencing when applied to the highly repetitive regions of the Y chromosome [21]. Despite this shortcoming, this analysis should give an indication of the magnitude of the change for the Y chromosome mutation rate. We find that no reasonable change in male generation time could account for the apparent significant increase in mutation rate since the split with Neandertals, as differences in generation time could only explain up to a 15% increase in mutation rate (Figure 2b). Further, we would expect generation time changes to affect all haplogroups in a population, rather than specific lineages as observed in the data. Therefore, while changes in paternal generation time may explain subtle differences in branch lengths in the Y chromosome phylogeny, they are unlikely to explain the full extent of the difference we observe.

### Exploring alternative hypotheses for the origin of branch length variation

Since neither biological nor demographic factors fully explain the branch length variation in the human Y chromosome phylogeny, we attempted to determine the extent to which reference bias could cause us to undercount mutations in Y chromosomes that are diverged from the reference used for alignment.

### Mapping to the chimpanzee reference Y chromosome

We first tested whether the use of an outgroup Y chromosome (that of the chimpanzee) as a reference can correct for the branch length variation in the human Y chromosome phylogeny. Although this strategy should result in a significant level of reference bias for all human Y chromosomes, we expect this bias to be the same across all lineages, as all are equally diverged from the chimpanzee Y chromosome reference. As we obtained many heterozygous genotype calls with the alignments to the chimpanzee reference (suggesting a high frequency of mapping errors) we concluded that human Y chromosomes are too diverged from the chimpanzee Y chromosome to allow us to accurately call genotypes (Table S3).

### Mapping to different human Y chromosome assemblies

Due to the highly repetitive nature of the human Y chromosome, more than half of the hg19 reference Y chromosome could not be assembled using short read sequencing methods. A fully contiguous telomere-to-telomere (T2T) Y chromosome assembly [11] (haplogroup J1a2b) allows us to test whether the use of a higher-quality reference influences the number of mutations we are able to detect by facilitating the alignment of more sequences in complex regions. A Y chromosome reference that is outside of the variation of most human haplogroups (haplogroup A0b [12]) provides an opportunity to test whether we can recover more mutations from the more basal lineages in the phylogeny.

To study the effect of the reference used for alignment and determine whether reference bias affects mapping to the Y chromosome, we aligned whole-genome sequence data from the individuals in our dataset for whom we had unmapped reads to three different reference genomes:

- The hg19 reference with the original hg19 Y chromosome which is almost entirely derived from a lineage that represents the R1b haplogroup (one that is within the variation of present-day non-African Y chromosomes)
- The hg19 reference with a long-read assembled Y chromosome representing the A0b lineage [12] (one that is outside of the variation of most modern human Y chromosome lineages, with only the A00 lineage as a more basal outgroup)
- The hg19 reference with the long-read assembled T2T Y chromosome representing the J1a haplogroup [11] (which, like the hg19 Y chromosome, is within the variation of present-day non-African Y chromosomes)

For each of the three reference genomes, we processed our dataset using the same pipeline, called genotypes, and compared the relative branch length differences with respect to a present-day non-African Y chromosome (Figure 3a). We find that the T2T reference has little effect on the branch length variation and we observe similar branch length differences for the two references that are within the variation of non-African Y chromosomes. This shows that mutations are not missed due to the quality of the reference assembly. By contrast, the A0b reference enables the detection of more mutations on the A0a1 lineage, but does not resolve the issue for the other A haplogroups that do not fall on the A0 lineage. We also miss mutations on the non-African lineages that are now highly diverged from the A0b reference (Figure 3b). Taken together, these results show that reference bias does affect the alignment of sequences from human Y chromosomes and that this bias will always be present for some lineages in the phylogeny depending on the reference used for alignment.

**Fig. 3:**
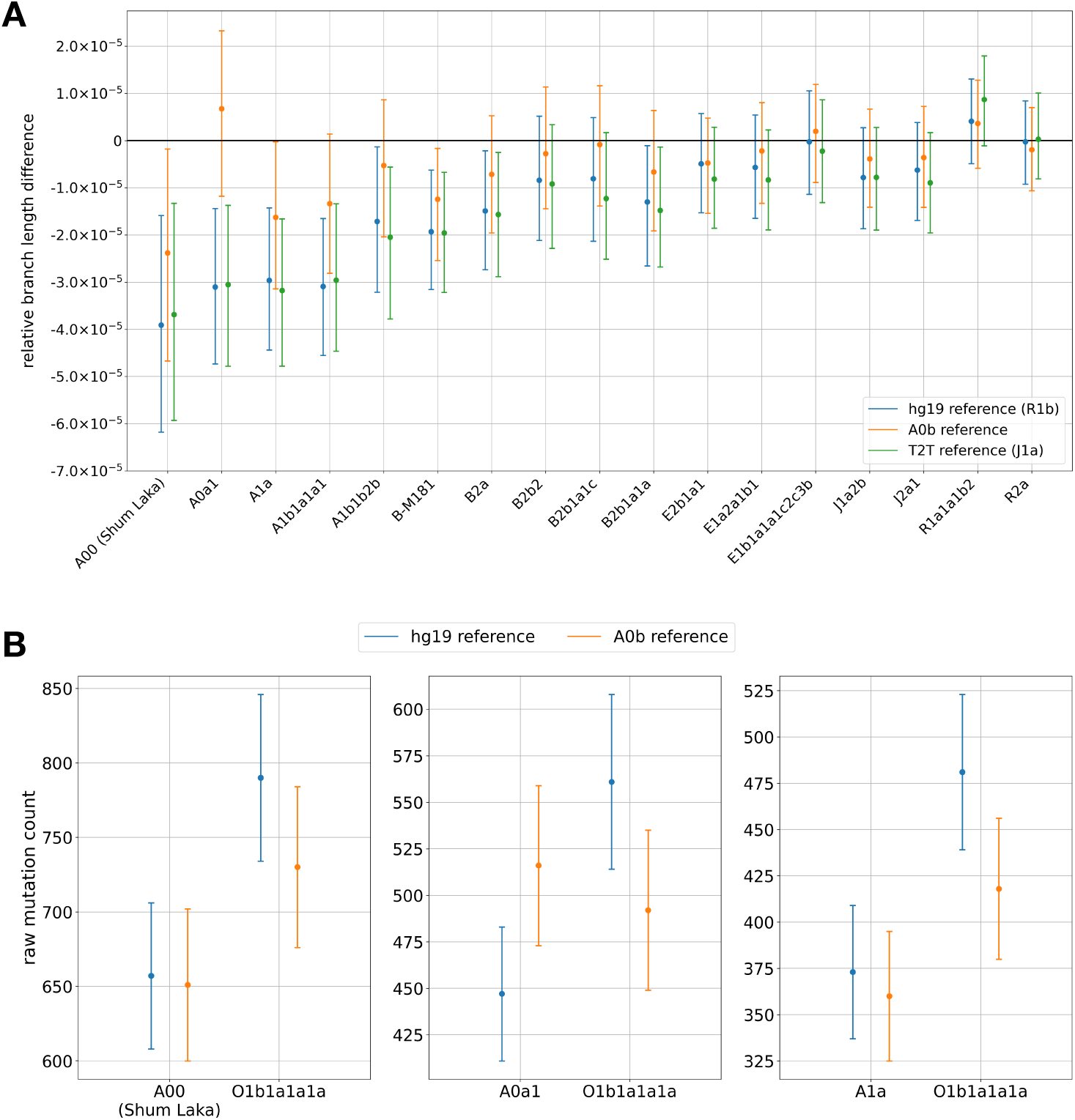
(a) Relative branch length differences compared to a present-day non-African lineage (O1b1a1a1a) using variants called from alignments to three different reference genomes (shown with different colours). The error bars correspond to 95% CIs computed by resampling branch lengths from a Poisson distribution. (b) Counts of private mutations for three present-day A lineages (A00, A0a1, and A1a, as indicated in each panel) since their last common ancestor with a present-day non-African lineage (O1b1a1a1a) using variants called from alignments to two different reference genomes (shown with different colours). The results with the alignments to the T2T reference genome are not shown, for simplicity, and are similar to those obtained with the hg19 reference.

### An approach to limit the effect of reference bias

As the reference used for alignment cannot resolve the effect of reference bias for all Y chromosome haplogroups, we then investigated whether specific regions of the hg19 Y chromosome are correlated with a reduction in the ability to call private mutations in divergent Y chromosomes. We used sequence divergence between the hg19 Y chromosome assembly and the chimpanzee reference (panTro6) Y chromosome assembly to identify the faster-evolving regions of the human Y chromosome where the alignment of short read data may be most affected by reference bias. We used a sliding window approach to compute the sequence divergence for each uniquely mappable base of the hg19 Y chromosome based on the number of mismatches compared to the panTro6 Y chromosome. For three present-day haplogroups that are affected by reference bias (A00, A0a1 and B-M181), we computed the mean mutation count difference with respect to all of the haplogroups that do not show significant branch length differences (i.e. all non-African haplogroups along with the E haplogroups) for progressively increasing sequence divergence filters (Figure 4). This allowed us to identify the sequence divergence cutoff that minimises the branch length difference between the lineages that are most diverged from the reference and those that are closer to the reference. As the non-recombining part of the Y chromosome is a single locus, only one phylogeny exists for the human Y chromosome. For this reason, removing regions of the Y chromosome does not introduce biases if we use the appropriate mutation rate in subsequent analyses.

**Fig. 4:**
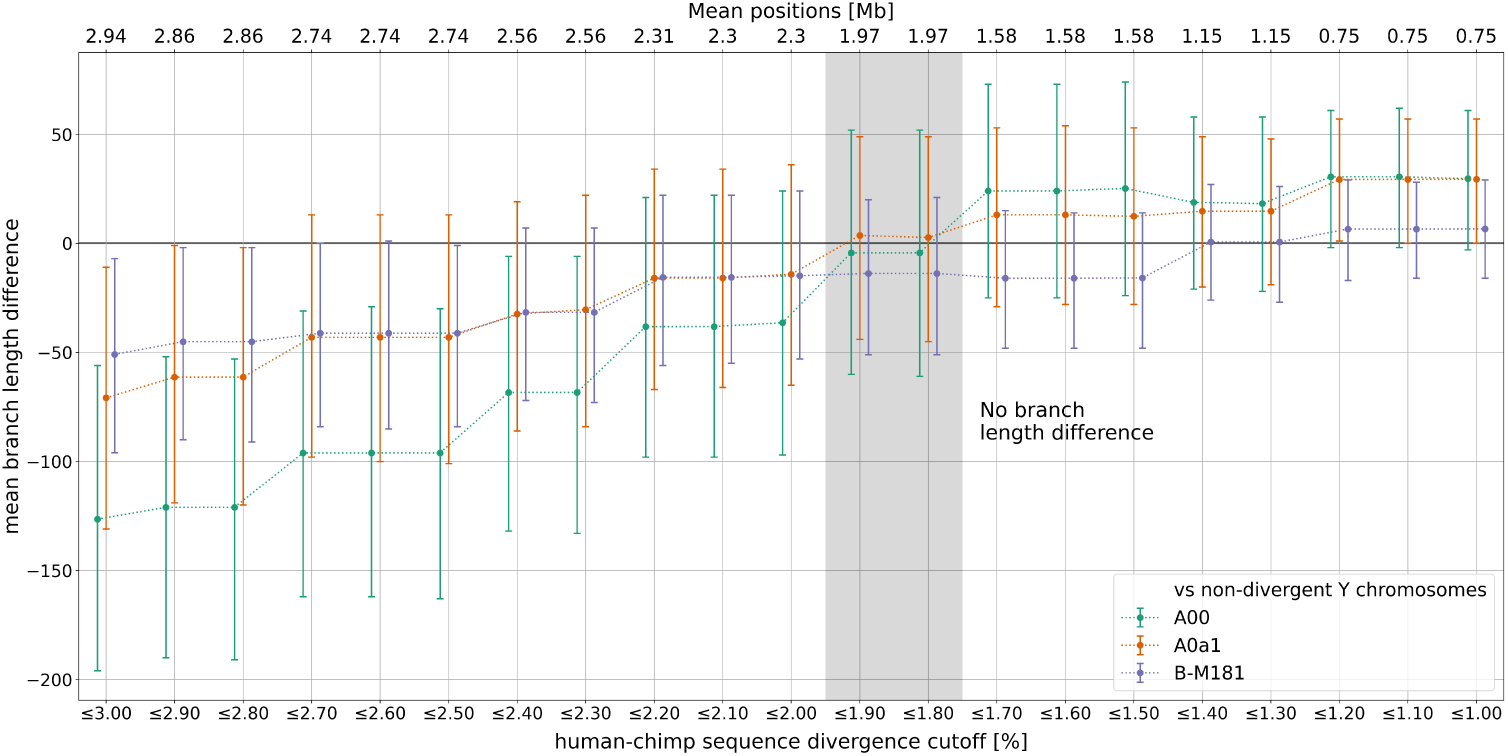
Comparing the number of mutations on divergent Y chromosomes to non-divergent Y chromosomes for different maximum human-chimp sequence divergence filters (lower x-axis). The upper x-axis represents the average number of positions (in Mb) used for each comparison. The colours correspond to the three divergent Y chromosomes (A00, A0a1 and B-M181). The shaded area indicates the divergence filters that minimise the branch length variation while maximising the proportion of the Y chromosome available for further analysis. The error bars denote 95% CIs computed by resampling branch lengths from a Poisson distribution.

There is a tradeoff, however, between filtering the regions of the Y chromosome where mapping is more difficult while simultaneously retaining a sufficient number of positions for further phylogenetic analysis, and we aimed to retain as many of the uniquely mappable positions of the human Y chromosome as possible. The branch length variation is minimised, and the number of positions maximised, using a human-chimp sequence divergence cutoff of 1.9% which retains around 2 Mb (megabases) of the uniquely mappable *∼*4.6 Mb of the hg19 Y chromosome.

We also looked at human-chimp sequence divergence for the complete nuclear genome to determine whether reference bias could have a similar impact on analyses of the autosomes or the X chromosome (Table S4). The Y chromosome stands out for both the proportion of the chromosome that can be aligned and the divergence from the chimpanzee reference, suggesting that reference bias is largest on the Y chromosome. This probably results from the faster evolving nature of the Y chromosome compared to the autosomes [14].

We then analysed the regions of the Y chromosome that are removed by the human-chimp divergence filter. About half of the non-recombining part of the human Y chromosome consists of highly repetitive heterochromatin and the remaining euchromatin comprises three sequence classes: the X-degenerate regions, the X-transposed regions, and the ampliconic regions, which primarily consist of palindromic repeats. We find that the majority of the uniquely mappable positions in the ampliconic and X-transposed regions are removed by the divergence filter, whereas more than half of the X-degenerate positions are retained (Table 1). We investigated whether the divergence filter is required in the X-degenerate regions of the Y chromosome by using only these regions when testing for the presence of branch length variation. We still observe shortened branches for the Y chromosome lineages that are most diverged from the reference (Figure S2) showing that reference bias also affects the X-degenerate regions and that the divergence filter is necessary for unbiased analyses of the human Y chromosome phylogeny.

**Table 1:** Uniquely mappable positions removed by the human-chimp divergence filter for each euchromatic sequence class of the Y chromosome.

| Sequence class | Total mappable positions | Positions retained | % Removed |
| --- | --- | --- | --- |
| X-degenerate | 3,701,385 | 2,151,164 | 41 |
| X-transposed | 27,214 | 0 | 100 |
| Ampliconic | 834,298 | 38,196 | 95 |
| Others | 39,452 | 11,458 | 71 |
| Total | 4,602,349 | 2,200,818 | 52 |

We hypothesized that regions masked by the divergence filter would show evidence for an increased rate of mapping errors. To assess this we identified heterozygous sites called by snpAD – a diploid genotype caller – on the haploid Y chromosome, and calculated the percentage change in the number of heterozygous calls for each individual after applying the human-chimp sequence divergence filter. We found that a large proportion (between 7 to 92%) of mapping errors occur in the divergent regions of the Y chromosome (Table S5), indicating the increased difficulty of correctly aligning reads in quickly-evolving regions.

### Application to estimates of the modern human and the modern human-Neandertal Y chromosome TMRCAs

After minimising the effect of reference bias in the Y chromosome phylogeny, we sought to recalibrate both the modern human Y chromosome TMRCA (using the A00 Y chromosome to represent the most basal modern human lineage) and the modern human-Neandertal Y chromosome TMRCA using the portion of the human Y chromosome with minimal reference bias. We used two different methods to determine the Y chromosome TMRCAs: 1) the method defined in [1] (with the lineages used shown in Figure 5) which computes the modern human-Neandertal TMRCA by scaling the modern human TMRCA based on modern human-Neandertal divergence and 2) Bayesian phylogenetic reconstruction as implemented in the software package BEAST2 (v2.6.7) [23].

**Fig. 5:**
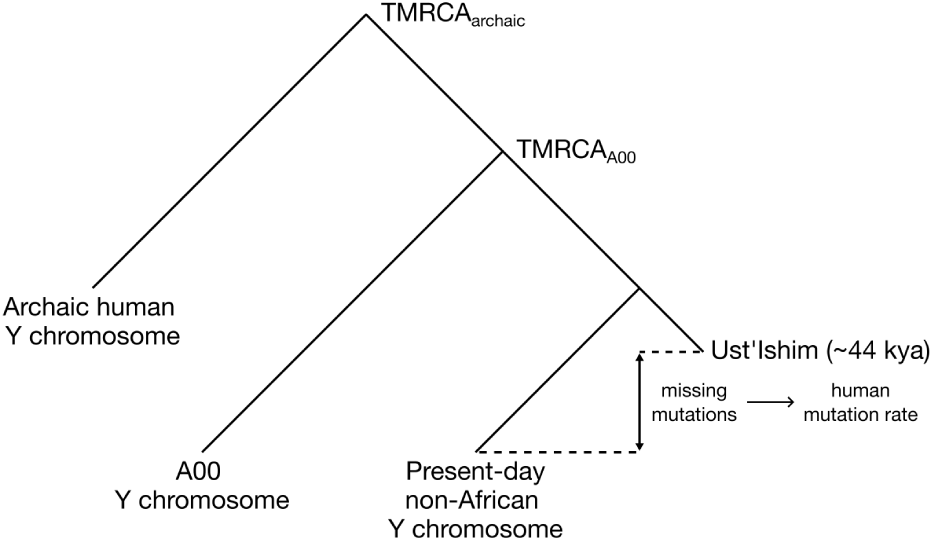
Tree depicting the lineages used to estimate the TMRCA of all modern humans and the TMRCA of modern and archaic humans. The lineage of the radiocarbon dated ancient human, *Ust’Ishim*, used to calculate the mutation rate, is also shown.

We first calculated the Y chromosome mutation rate for those regions of the Y chromosome that pass the divergence filter (Table 2). The mutation rate in these regions is slightly lower than the mutation rate computed over the entire uniquely mappable Y chromosome (Table 2) and the rates reported elsewhere (Table S6). This is expected as more divergent, faster-evolving regions of the Y chromosome are removed by the divergence filter. In all subsequent analyses, we used this recalibrated mutation rate to account for the effect of the divergence filter.

**Table 2:** Comparing the mutation rate estimates (in units of mutations/bp/year) for the uniquely mappable part of the Y chromosome and for the uniquely mappable, divergence-filtered Y chromosome (maximum 1.9% human-chimp sequence divergence). The method used to estimate the mutation rate (as described in the main text) is indicated. HPD - highest posterior density.

| Method | Unfiltered Y chromosome | Divergence-filtered Y chromosome |
| --- | --- | --- |
| Branch shortening | $7.22 \times 10^{-10}$ | $6.66 \times 10^{-10}$ |
| | 95% CI $4.84 \times 10^{-10}$ to $9.85 \times 10^{-10}$ | 95% CI $4.22 \times 10^{-10}$ to $9.57 \times 10^{-10}$ |
| BEAST | $7.28 \times 10^{-10}$ | $6.80 \times 10^{-10}$ |
| | 95% HPD $6.68 \times 10^{-10}$ to $7.86 \times 10^{-10}$ | 95% HPD $6.06 \times 10^{-10}$ to $7.54 \times 10^{-10}$ |

We then computed the modern human and the modern human-Neandertal TMR-CAs, and compared the TMRCAs estimated with all of the uniquely mappable sites to those estimated with only sites that pass the divergence filter (Figure 6). When the unfiltered Y chromosome is used, the choice of the modern human branch used for comparison has an effect on the TMRCA calculation with the difference in length between the A00 branch and the non-African branches resulting in inconsistent estimates. By applying the divergence filter, the effect of reference bias is minimised and the choice of modern human branch then has no effect on the TMRCA estimates.

**Fig. 6:**
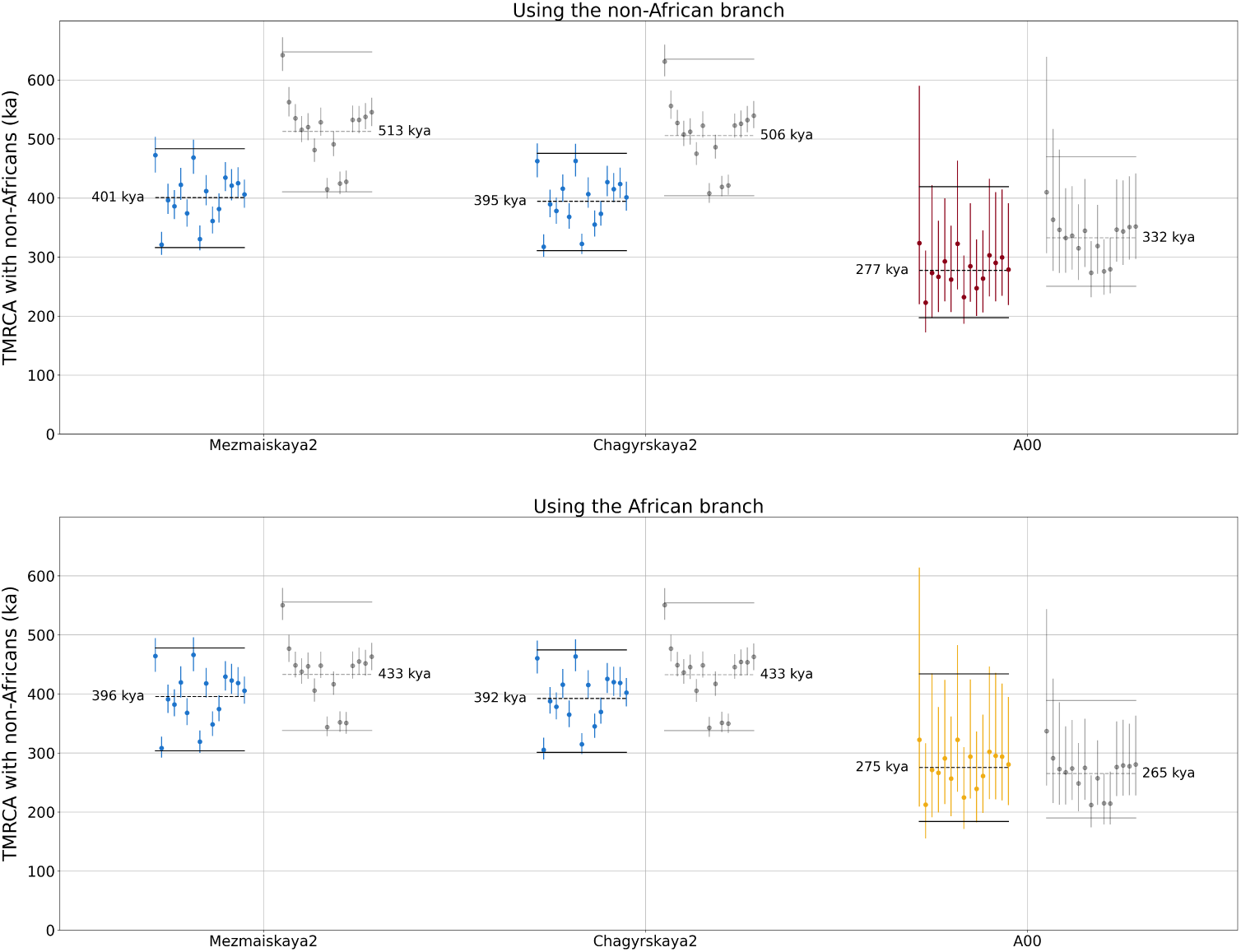
TMRCA estimates between the Y chromosomes on the x-axis and 16 non-African Y chromosomes. Each dot represents the TMRCA with one non-African Y chromosome. The top plot shows the TMRCAs estimated by measuring the non-African branch, while the bottom plot shows the TMRCAs estimated by measuring the A00 branch. The dots in colour represent the TMRCA estimates based on the the filtered Y, while the dots in grey represent the TMRCAs estimated using the unfiltered Y chromosome. The vertical lines denote 95% CIs computed by resampling branch lengths from a Poisson distribution. The dashed horizontal lines represent the mean TMRCAs computed over all non-African Y chromosomes and the solid horizontal lines show overall 95% CIs.

As we are able to calculate unbiased TMRCAs after masking divergent regions, we then used BEAST to reconstruct the Y chromosome phylogeny (see Methods) based on the sites that pass the divergence filter (Figure 7). BEAST estimates similar TMRCAs to those estimated with the method described above, although with narrower confidence intervals. We estimated that the modern human TMRCA is 268 ka (95% HPD 238 to 301 ka) and the modern human-Neandertal TMRCA is 387 ka (95% HPD 344 to 432 ka). Our recalibrated TMRCAs are slightly older than previous estimates of 254 ka (95% CI 192 to 307 ka) for the modern human TMRCA [7] and 370 ka (95% CI 326 to 420 ka) for the modern human-Neandertal TMRCA [1], but within their confidence intervals.

**Fig. 7:**
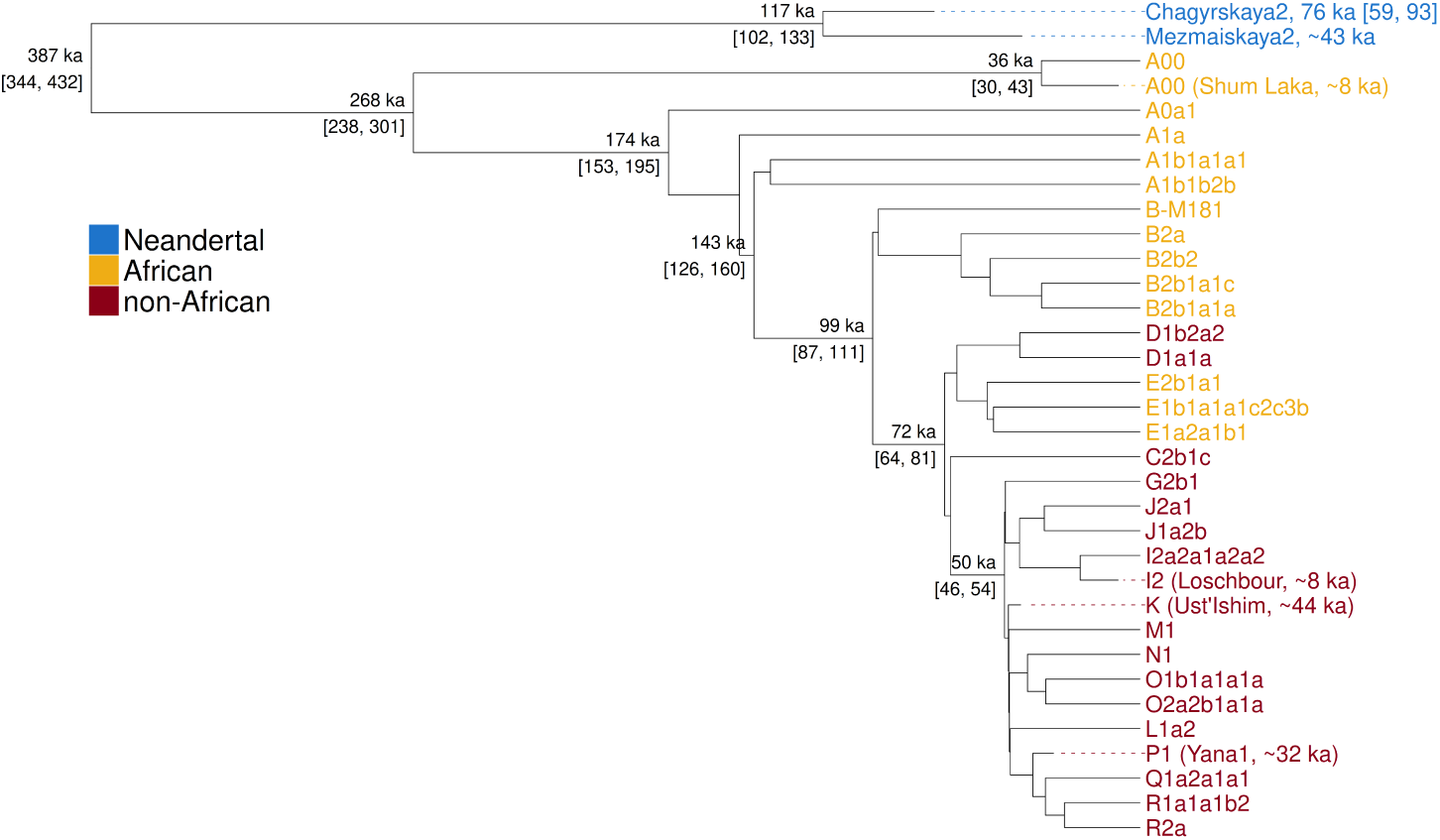
Y chromosome phylogeny reconstructed with BEAST. The TMRCA estimates, as well as the estimated age of *Chagyrskaya 2*, and their respective 95% HPD intervals are indicated on the tree. The ages of the other ancient samples were set to the estimated radiocarbon dates. The haplogroups from different populations are highlighted with different colours. The branches are to scale, in thousands of years (ka).

We then tested whether we can use BEAST to estimate the age of a shotgun sequenced ancient individual from Shum Laka, Cameroon dated to *∼*8 ka (7,970–7,800 calibrated years before present) who carried the A00 Y haplogroup using just the Y chromosome. For this analysis, we set a wide prior for the age of the ancient individual (-50,000 to 200,000) and did not include the present-day A00 individuals. Using the same parameters used for generating the full phylogeny above, we constructed one phylogeny with the filtered Y chromosome and one with the unfiltered Y chromosome (Figure 8). We find that if we use the unfiltered Y chromosome to construct the phylogeny, we get an age-estimate for the ancient A00 individual that is about 30 thousand years older than the radiocarbon date due to the reference bias that leads to an underestimate of the number of mutations on his branch. If we use the filtered Y chromosome regions, we obtain an age estimate of 13 ka (95% CI 43 to 0) which is reasonably close to the radiocarbon date (albeit with wide confidence intervals) indicating that the phylogeny is unbiased.

**Fig. 8:**
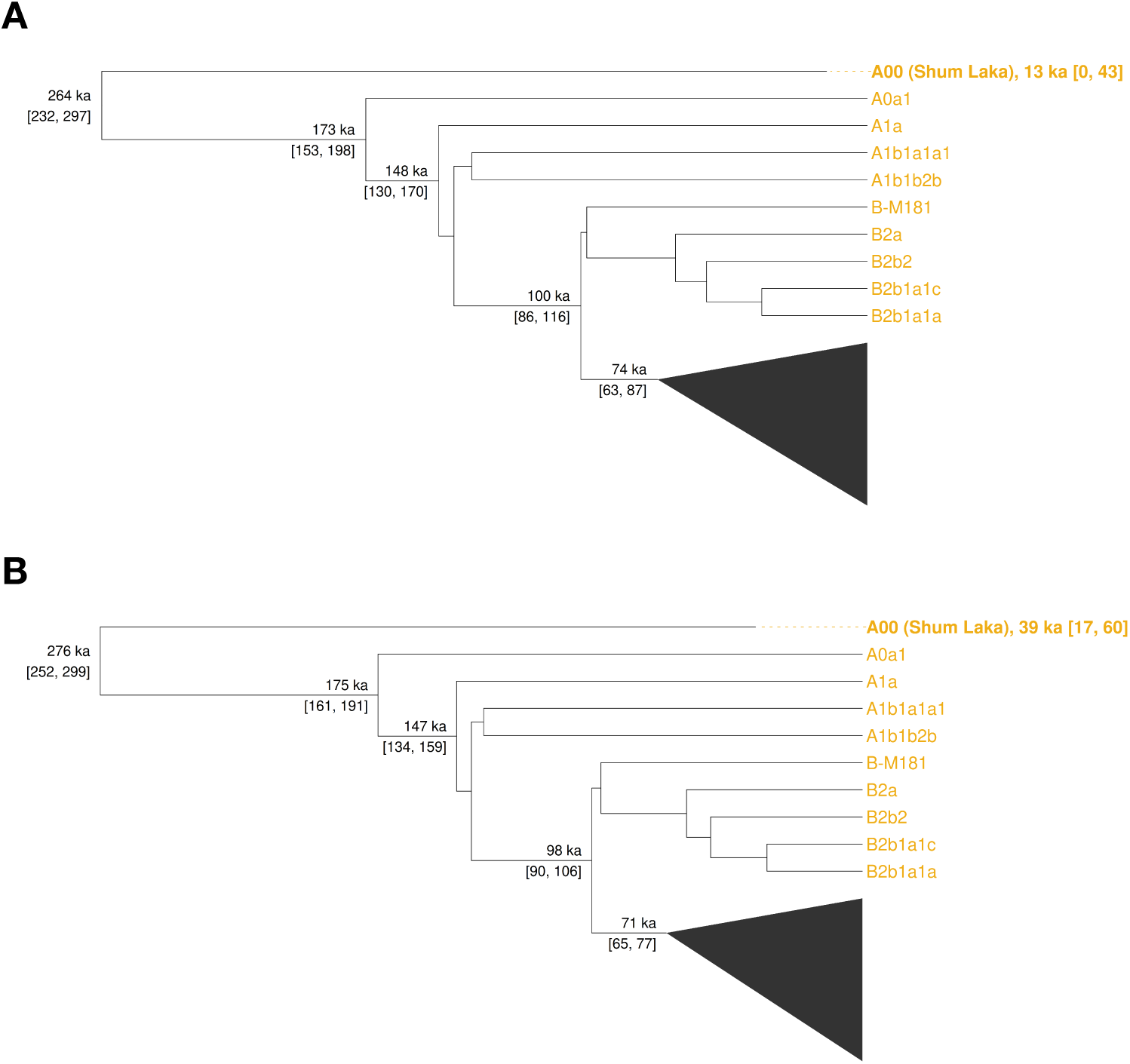
Estimating the age of the radiocarbon dated A00 individual (radiocarbon date *∼*8 ka) using molecular dating of the Y chromosome with BEAST. Phylogenies based on the filtered (a) and the unfiltered (b) Y chromosome are shown. The branches corresponding to 21 Y chromosomes included in the analysis (Table S1) were collapsed. The TMRCA estimates and their 95% HPD intervals are indicated on the tree. The age of the A00 individual, along with its associated 95% HPD interval, is indicated in the branch label.

We also re-estimated an age for the *Chagyrskaya 2* Neandertal — previously dated to *∼*64 ka (95% CI 48-83) [13]. Our estimate of 76 ka (95% CI 59-93) is consistent with a Neandertal occupation of Chagyrskaya Cave between 92 ka and 49 ka, and close to the genetic age of *Chagyrskaya 8* (*∼*80 ka), another individual from the same cave, whose age was estimated from a high-coverage genome [24].

In summary, although the re-estimated TMRCAs are not significantly different from previous estimates, these analyses show that the divergence filtering is necessary to obtain Y chromosome-based age estimates for ancient individuals with Y chromosomes that are diverged from the human reference.

## Discussion

We show here that variation in Y chromosome branch lengths is at least in part explained by biases introduced by alignment to a single Y reference genome representing a single haplogroup. Although we do not observe any significant haplogroup-specific variation after accounting for the effect of reference bias (Figure S3 and Figure S4), a larger Y chromosome dataset with multiple samples representing specific lineages in populations of interest may allow for the detection of subtle branch length differences that result from changes in mutation rate or generation time.

The Y chromosome is more affected than the autosomes by this reference bias due to its highly repetitive nature and extensive structural variation. We show that restricting analyses of human Y chromosomes to regions with a human-chimpanzee divergence of *≤*1.9% allows accurate estimation of branch lengths and, therefore, unbiased estimates of the human Y chromosome phylogeny. However, doing so reduces the (already limited) amount of the Y chromosome that is accessible for analysis by more than half, leading to wider confidence intervals in both TMRCA and age estimates. For analyses of Y chromosomes in specific populations, it may be possible to recover more of the Y sequence by alignment to the closest high-quality Y chromosome assembly [12].

In contrast to using a single linear reference genome, approaches that incorporate Y chromosome haplotype diversity in the reference to which sequence reads are aligned, known as ‘pan-genomes’, have been shown to reduce the effect of reference bias when used as a reference for short-read mapping [25, 26] and could potentially maximise the amount of the Y chromosome that is accessible for comparison. The repetitive nature of the Y chromosome, along with its extensive structural variation, makes it a particularly good target for a pangenome-based approach. A number of high-quality Y chromosome assemblies that could be used to construct a pangenome reference [12] are already available, although methods to incorporate ancient Y chromosome variation into pangenome graphs have yet to be explored and are an exciting direction for future research.

We expect our divergence-filtering approach to be particularly useful for analyses of archaic human (i.e. Neandertal and Denisovan) Y chromosomes that are substantially diverged from the human reference, and for which the construction of pangenome graphs remains challenging. These Y chromosomes will be most affected by reference bias, so methods that rely on comparing the number of private mutations on archaic Y chromosomes to those of present-day human Y chromosomes will underestimate the archaic branch length if they do not account for this bias.

## Materials and Methods

### Dataset

The dataset includes high-coverage Y chromosome data from present-day and ancient modern humans as well as from Neandertals (Table S1). For the present day humans, a subset of individuals from the Human Genome Diversity Project (HGDP) [18] dataset were selected to represent the Y chromosome phylogeny with a particular focus toward individuals with Y chromosomes that are highly diverged from the human reference Y chromosome. We also included two individuals (carrying Y chromosomes representing the A haplogroup) from the high-coverage resequencing of the 1000 Genomes Project (1KGP) [19].

The dataset also contains Y chromosomes from two closely related individuals with the A00 haplogroup [7]. Due to the relatively low coverage of the sequence data for these two A00 Y chromosomes, we followed the approach of the original publication and merged the two A00 Y chromosomes into a single BAM file.

To calibrate the Y chromosome phylogeny, we included high-coverage, shotgun-sequenced Y chromosome data from the following radiocarbon dated ancient modern humans: *Ust’Ishim* [14], *Yana1* [17], *Loschbour* [16] and an individual from Shum Laka with the A00 haplogroup [15].

We also included two high-coverage Neandertal Y chromosomes that were obtained by enrichment-capture sequencing: *Mezmaiskaya2* [1] and *Chagyrskaya2* [13].

### Data Processing

Sequences were aligned using BWA-MEM (v0.7.17) [27] for present-day individuals and BWA-ALN (with ancient parameters — “-n 0.01 -o 2 -l 16500”) for ancient individuals. For the present-day HGDP and 1KGP individuals, adapter sequences were identified and flagged using MarkIlluminaAdapters from Picard tools (v2.18.29) before alignment. For the ancient samples, we used the merged and adapted trimmed sequences as published (Table S1).

The aligned BAM files for each individual were filtered with samtools [28] to retain reads at least 35 bp in length and with a minimum mapping quality of 25, and PCR duplicates were removed using bam-rmdup (v0.2, https://github.com/mpieva/biohazard-tools/). We also removed sequences with indels as we observed an increased frequency of genotyping errors around indels.

We restricted our analysis to uniquely mappable regions of the Y chromosome so as to minimise errors caused by ambiguous alignment of short reads. To exclude non-unique regions of the hg19 Y chromosome, we used the map35_100 filter from [29] which retains only positions where all 35mers overlapping that position do not map to any other position in the genome allowing up to one mismatch. We also applied the capture_full filter from [1], which includes masks of tandem repeats identified by Tandem Repeat Finder [30].

Genotypes were called with snpAD [31] – a genotyper that accounts for ancient DNA damage – using bases with a minimum base quality of 30. For consistency, snpAD was used to call genotypes for both ancient and modern individuals, however, position-specific error rates were estimated independently for each sample. As snpAD is designed to call diploid genotypes, heterozygous genotype calls were converted to homozygous genotypes choosing the lowest homozygous posterior probability as calculated by snpAD. Finally, following [29], genotypes were filtered based on maximum GC-corrected coverage cutoffs for each individual (see Availability of data and materials) in order to remove potential duplications or regions with spurious alignments of microbial sequences.

## Methods to Estimate Branch Shortening

### Branch Length Difference

To calculate the difference in branch length with respect to a reference individual, we counted the number of derived mutations on each branch (using the chimpanzee genome to infer the ancestral state) with the *BranchShortening.get_mutations_vs_individual* function from https://github.com/yanivsw/y_ chr_utils. The UCSC whole genome alignments between the human reference genome (hg19) and the chimpanzee reference genome (panTro6) were used to determine ancestral genotypes.

### Estimating the Human Y chromosome Mutation Rate

A Y chromosome from an ancient human individual with an accurately estimated age can be used to determine the human Y chromosome mutation rate. The *Ust’Ishim* individual is estimated, based on radiocarbon dating, to have lived around 44 thousand years ago (45,930-42,904 calibrated years before present; [14]). We used the high-coverage genome sequence of *Ust’Ishim* to determine the number of derived mutations on the branch leading to *Ust’Ishim*, and compared this to an individual living today. Normalising the number of mutations accumulated per year by the total number of positions used for the comparison results in the Y chromosome mutation rate in units of mutations per position per year.

### Estimating the Mutation Rate on a Shortened Branch

To estimate the mutation rate that would result in a shortened branch when comparing two Y chromosome lineages, we first calculate the expected branch length difference using the known ages of the two individuals and the expected mutation rate. We then calculate the proportion of missing mutations using the observed branch length difference and adjust the expected mutation rate, based on this proportion, to estimate the mutation rate on the shorter branch.

### Estimating TMRCAs of Modern and Archaic Human Y Chromosomes

We used the method described in [1] to estimate the TMRCAs of modern and archaic Y chromosomes. Briefly, the method first estimates the modern human TMRCA by counting the differences between a present-day African Y chromosome and a present-day non-African Y chromosome and converting this count to time using a mutation rate. The archaic TMRCA is then computed by scaling the modern human TMRCA based on the fold increase in divergence between one of the present-day Y chromosomes and the archaic Y chromosome relative to the divergence between the two present-day Y chromosomes.

### Mapping to the Chimpanzee Reference Y Chromosome

We replaced the hg19 Y chromosome with the chimpanzee reference (panTro6) Y chromosome (hg19_panTro6_Y) and aligned all sequence reads from two ancient genomes (*Ust’Ishim* and *Shum Laka*) to this reference. To conduct a fair comparison, we downsampled the aligned sequences of *Ust’Ishim* to the same coverage distribution and read length distribution of the aligned sequences of *Shum Laka*. We then called genotypes with the same pipeline that we use to call genotypes from the sequences aligned to the human hg19 reference.

### Estimating the Percentage Change in Mutation Rate in Response to a Change in Generation Time

To estimate the relative change in mutation rate corresponding to a change in generation time, we used the relationship between paternal age (*a*) and the autosomal germline mutation rate (*µ*, in units of male-transmitted mutations per generation) defined in [32] as

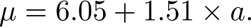

To account for the fact that a decrease in the average paternal age can result in an increase in the number of mutations over time due to an increase in the number of generations, we define the relative change in mutation rate (if the paternal age changed from *a_initial_* to *a_current_*) as

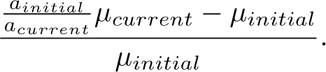

### Processing of Long-Read Assembled Y Chromosomes

We replaced the Y chromosome of the hg19 reference genome with the A0b Y chromosome (hg19_A0b_Y) from [12] or with the T2T Y chromosome (hg19_T2T_Y) from [11]. We then generated new map35_100 filters and ran Tandem Repeat Finder [30] on both the hg19_A0b_Y genome and the hg19_T2T_Y genome to identify the uniquely mappable regions for each genome. To call genotypes, we used the same processing pipeline as the one used for hg19.

To identify ancestral alleles, we used LASTZ [33] to align the chimpanzee reference genome (panTro6) to both the hg19_A0b_Y genome and the hg19_T2T_Y genome using the UCSC LASTZ alignment parameters for human-chimp alignments. The alignments were then chained with axtChain using the UCSC chaining parameters and the high-scoring (*>* 2000) alignments were extracted from which we ‘called’ panTro6 genotypes.

### Human-chimp Sequence Divergence

To compute human-chimp sequence divergence for each base of the hg19 Y chromosome, we extracted the Y chromosome alignments from the UCSC hg19-panTro6 pairwise alignment file (hg19.panTro6.synNet.maf.gz) and used the *maf_to_pairwise_identity.py* script from [34] to determine the alignment state (i.e. match, mismatch, insertion, deletion, no alignment) for each position in the hg19 Y chromosome. We then used a 400 bp sliding window centred on each position to compute human-chimp sequence divergence as

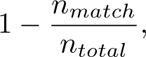

where *n_match_* is the number of matching bases and *n_total_* is the total number of aligned bases in each window. Due to the fragmented nature of the uniquely mappable fraction of the human Y chromosome, we additionally required a minimum number of human-chimp aligned bases (200 bp) to compute the sequence divergence for a position.

### Bayesian Phylogenetic Analysis

We used BEAST2 (v2.6.7) [23] to reconstruct the Y chromosome phylogeny and estimate the TMRCAs. As BEAST requires a FASTA file as input, the *VcfHandler.convert_vcf_dataframe_to_fasta_file* function from https://github. com/yanivsw/y_chr_utils was used to convert the Y chromosome VCF data to FASTA format. Only variable sites were used as input to the BEAST analysis in order to reduce the computational complexity of the phylogenetic reconstruction, although we defined the nucleotide composition of the invariant sites by editing the BEAST .xml file as was done in [7].

We used the BEAST2 bModelTest package [35] to estimate the most appropriate nucleotide substitution model for our data set. bModelTest averages over all substitution models while simultaneously reconstructing the phylogeny.

To select the tree model and the clock model that best fit our dataset, we estimated the marginal likelihood for each model combination using a path sampling approach as implemented in the BEAST2 MODEL_SELECTION package (Table S7).

To generate the full phylogeny using the best fitting models (the strict clock model together with the Bayesian Skyline tree model), we performed four independent MCMC runs of 55 million iterations with a 10 million iteration pre-burnin stage, sampling every 5,000 steps. We then combined the resulting log files with logcombiner using a 10% burnin for each run and generated a single best supported tree with TreeAnnotator.

## Declarations

### Ethics approval and consent to participate

All sequencing data used in this work was previously published.

### Consent for publication

Not applicable.

### Availability of data and materials

All sequence data used are publicly available (Table S1). The merged genotype files and filters used for the analysis are available from Zenodo (DOI:10.5281/zenodo.12635540) and all code is available from https://github.com/yanivsw/y_chr_reference_bias.

### Authors’ contributions

All authors contributed to the study conception and design. Data collection, programming and data analysis were performed by Y.S. The first draft of the manuscript was written by Y.S and S.P. All authors revised and approved the final manuscript.

## Acknowledgements

We thank Kay Prüfer and Leonardo N. M. Iasi for helpful discussions and Mark Stoneking and Eleni Seferidou for both helpful discussions and feedback on the manuscript.

## Supplementary Materials

### Bayesian Phylogenetic Analysis

The tip dates of modern samples were set to zero, while uniform priors were used to represent the ages of the ancient samples. For the radiocarbon dated ancient samples, the range of the prior was set as the 95% confidence interval based on the IntCal20 calibration curve and for *Chagyrskaya2* (who cannot be radiocarbon dated, but has been genetically dated to 64,000 years ago [13]) we set a wide uniform prior ranging from 40,000 to 120,000. We set the initial Y chromosome mutation rate to 7.34*×* 10*^−^*^10^ mutations/bp/year as calculated in [1]. In order to allow BEAST to effectively estimate the Y chromosome mutation rate, we set a wide uniform prior of 4*×*10*^−^*^10^ to 10*×*10*^−^*^10^.

The modern humans and the Neandertals were constrained as two different monophyletic groups in order to estimate their respective Y chromosome TMRCAs, and we estimated the modern human-Neandertal TMRCA by creating a third monophyletic group with only the chimpanzee as an outgroup.

To select the tree model and the clock model that best fit our dataset, we estimated the marginal likelihood for each model combination using a path sampling approach as implemented in the BEAST2 MODEL_SELECTION package. We tested the following model combinations:

- Strict clock, constant population size
- Relaxed log-normal clock, constant population size
- Strict clock, Coalescent Bayesian Skyline
- Relaxed log-normal clock, Coalescent Bayesian Skyline

For each model combination we used 50 path steps, an alpha parameter of 0.3 for the Beta distribution used to space out the steps, and a chain length of 10 million MCMC iterations for each step. The pre-burn-in stage was set at 5 million iterations and an 80% burn-in was used for each chain.

The two best supported model combinations were those with the Coalescent Bayesian Skyline tree model with slightly higher support for the relaxed clock model (Table S7). The support for the relaxed clock model over the strict clock model was not significant however (Bayes factor = 0.34), so we used the simpler model of the strict clock together with the Bayesian Skyline tree model to infer the various TMRCAs.

The nucleotide substitution model with the highest posterior probability as estimated by BEAST using the bModelTest package is the unnamed 123421 model. The nucleotide substitution rates are summarised in Table S8.

### Supplementary Figures

**Fig. S1:**
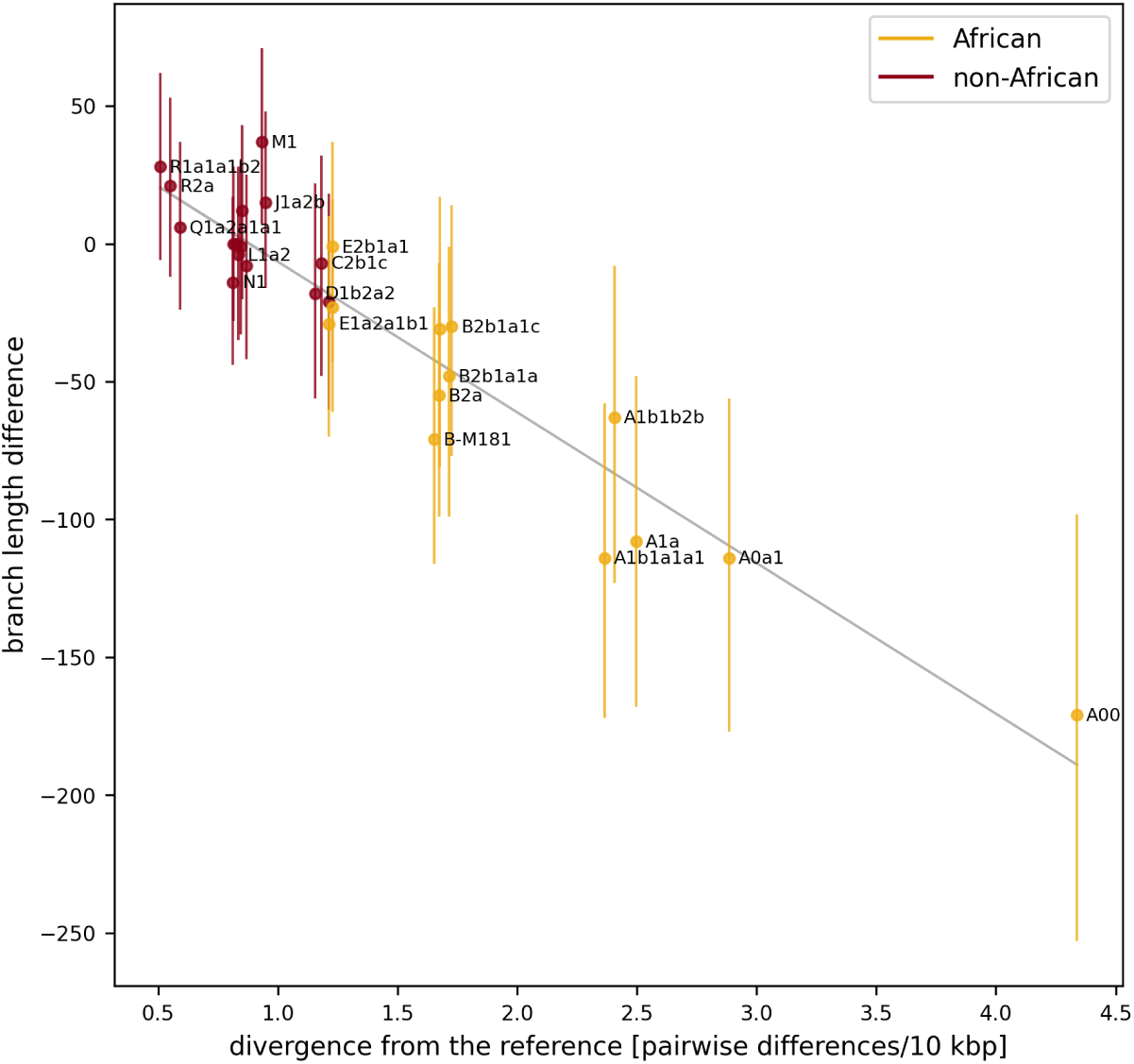
Branch length differences versus divergence from the reference for all present-day Y chromosomes. The black line represents the straight line that best fits the data with slope -54.66 mutations per pairwise differences per 10 kbp (kilobase pairs).

**Fig. S2:**
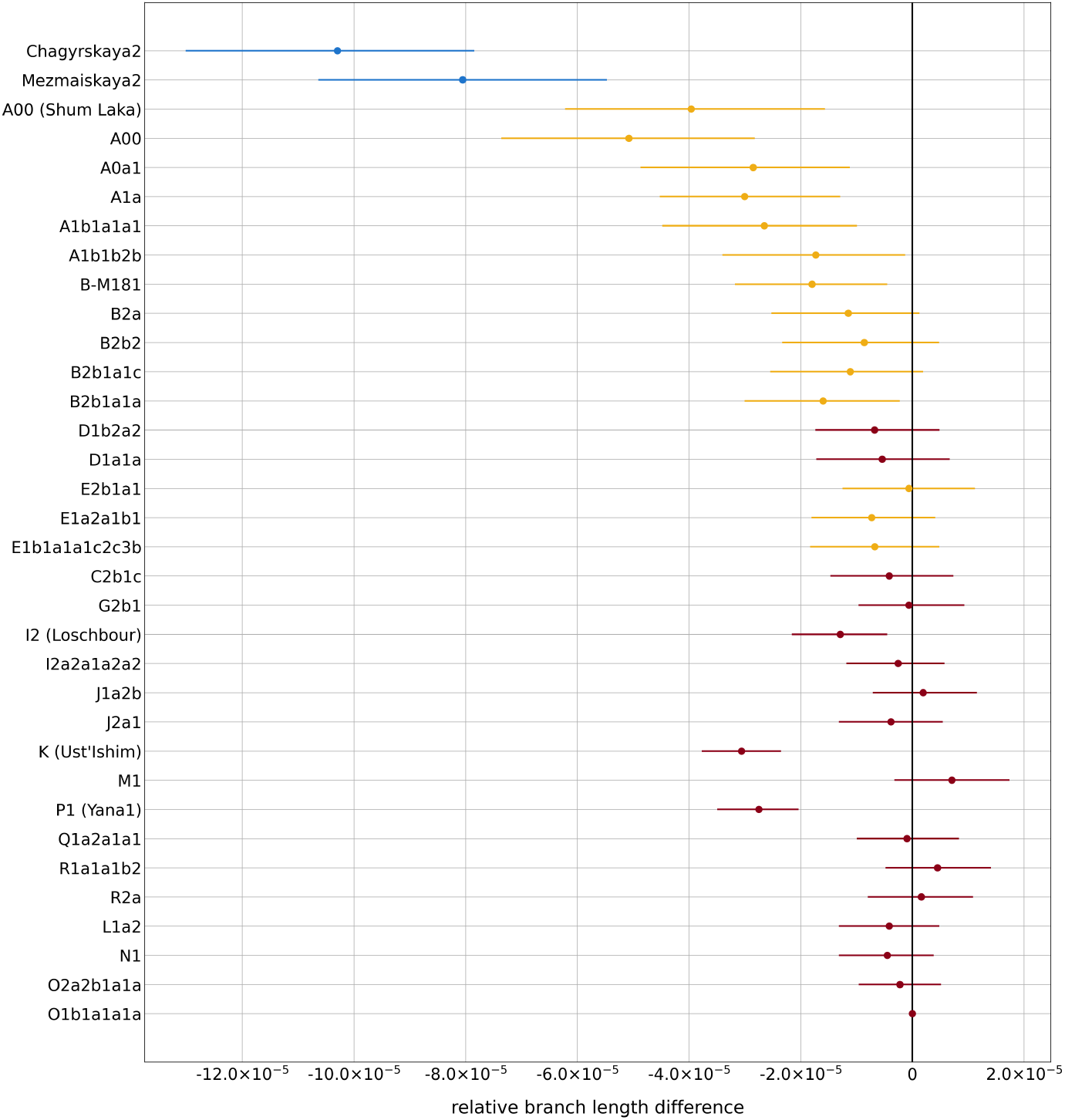
Relative branch length differences compared to a present-day non-African Y chromosome (O1b1a1a1a) based on the X-degenerate regions of the Y chromosome.

**Fig. S3:**
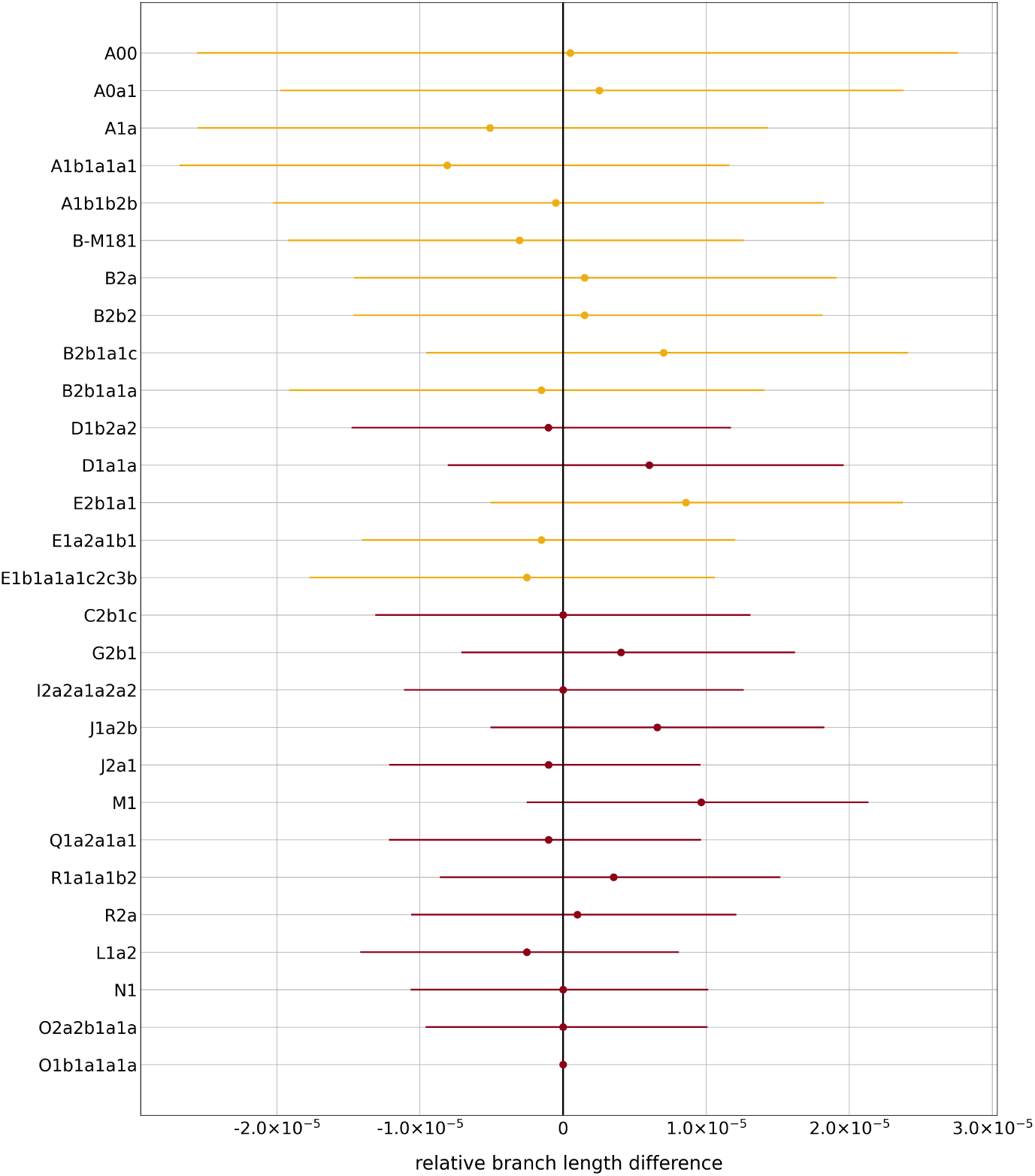
Relative branch length differences compared to a present-day non-African Y chromosome (O1b1a1a1a) for all present-day Y chromosomes after minimising the effect of reference bias.

**Fig. S4:**
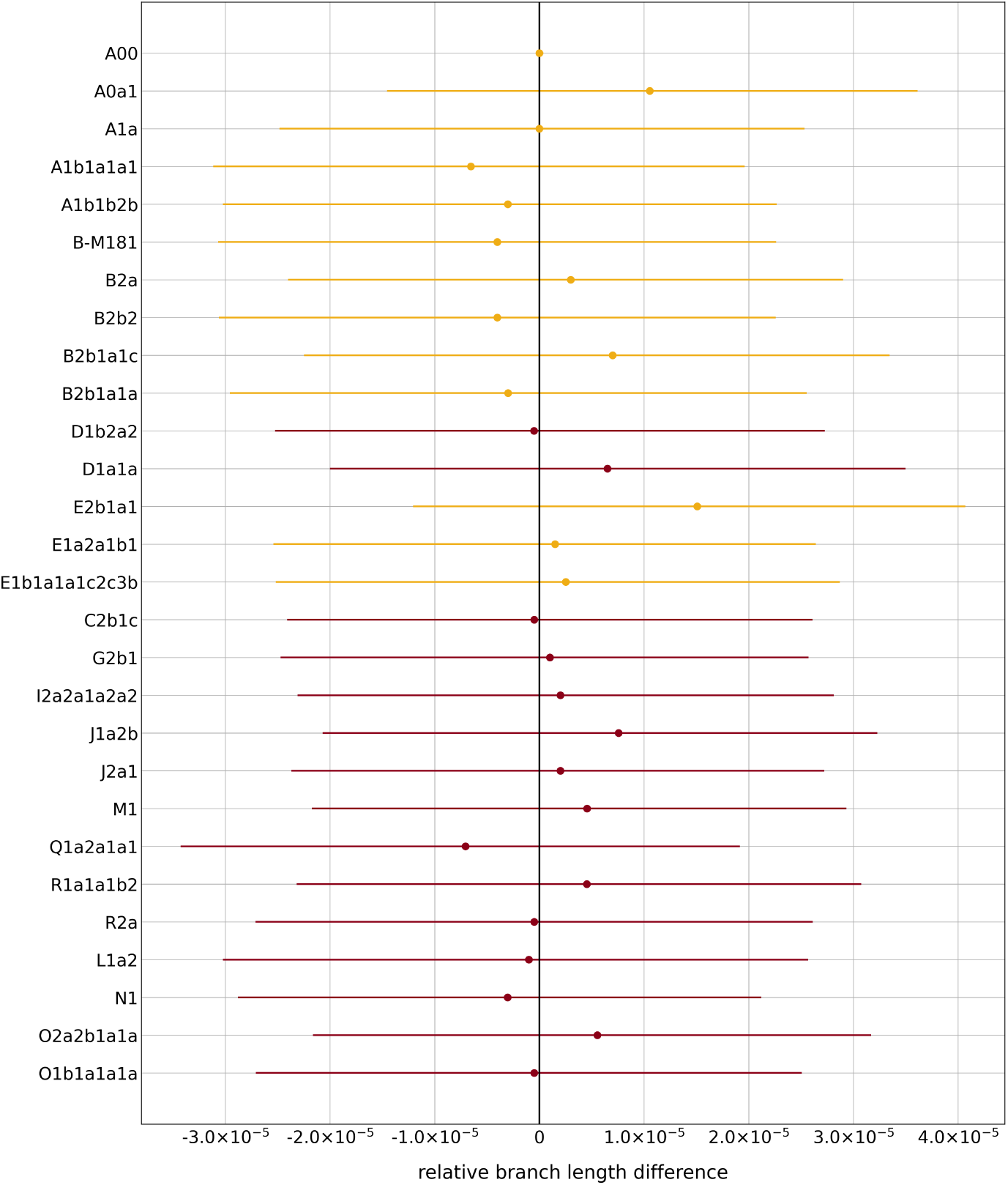
Relative branch length differences compared to A00 for all present-day Y chromosomes after minimising the effect of reference bias.

### Supplementary Tables

**Table S1:**
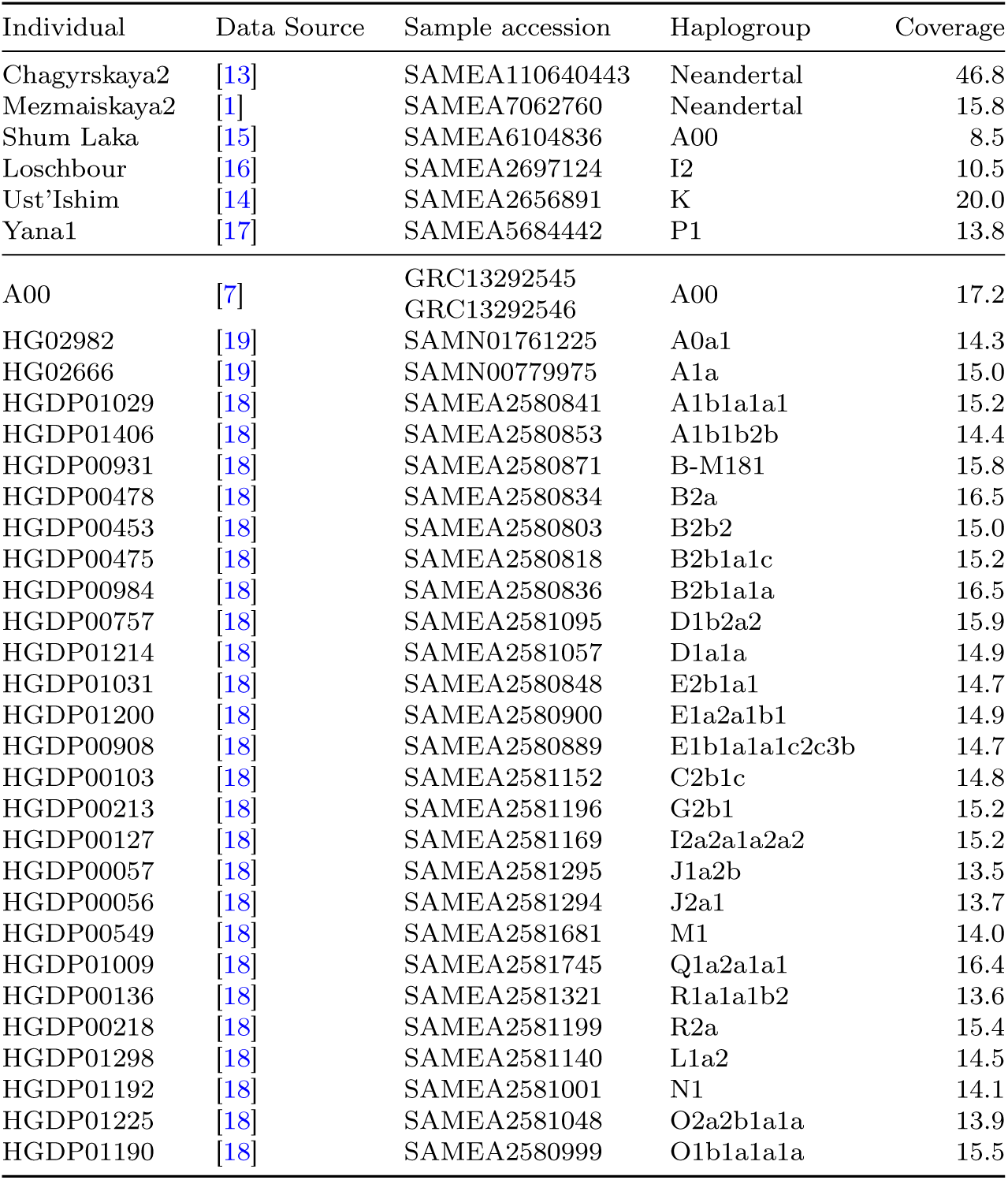
Dataset.

| Individual | Data Source | Sample accession | Haplogroup | Coverage |
| --- | --- | --- | --- | --- |
| Chagyrskaya2 | [13] | SAMEA110640443 | Neandertal | 46.8 |
| Mezmaiskaya2 | [1] | SAMEA7062760 | Neandertal | 15.8 |
| Shum Laka | [15] | SAMEA6104836 | A00 | 8.5 |
| Loschbour | [16] | SAMEA2697124 | I2 | 10.5 |
| Ust'Ishim | [14] | SAMEA2656891 | K | 20.0 |
| Yana1 | [17] | SAMEA5684442 | P1 | 13.8 |
| A00 | [7] | GRC13292545 | A00 | 17.2 |
|  |  | GRC13292546 |  |  |
| HG02982 | [19] | SAMN01761225 | A0a1 | 14.3 |
| HG02666 | [19] | SAMN00779975 | A1a | 15.0 |
| HGDP01029 | [18] | SAMEA2580841 | A1b1a1a1 | 15.2 |
| HGDP01406 | [18] | SAMEA2580853 | A1b1b2b | 14.4 |
| HGDP00931 | [18] | SAMEA2580871 | B-M181 | 15.8 |
| HGDP00478 | [18] | SAMEA2580834 | B2a | 16.5 |
| HGDP00453 | [18] | SAMEA2580803 | B2b2 | 15.0 |
| HGDP00475 | [18] | SAMEA2580818 | B2b1a1c | 15.2 |
| HGDP00984 | [18] | SAMEA2580836 | B2b1a1a | 16.5 |
| HGDP00757 | [18] | SAMEA2581095 | D1b2a2 | 15.9 |
| HGDP01214 | [18] | SAMEA2581057 | D1a1a | 14.9 |
| HGDP01031 | [18] | SAMEA2580848 | E2b1a1 | 14.7 |
| HGDP01200 | [18] | SAMEA2580900 | E1a2a1b1 | 14.9 |
| HGDP00908 | [18] | SAMEA2580889 | E1b1a1a1c2c3b | 14.7 |
| HGDP00103 | [18] | SAMEA2581152 | C2b1c | 14.8 |
| HGDP00213 | [18] | SAMEA2581196 | G2b1 | 15.2 |
| HGDP00127 | [18] | SAMEA2581169 | I2a2a1a2a2 | 15.2 |
| HGDP00057 | [18] | SAMEA2581295 | J1a2b | 13.5 |
| HGDP00056 | [18] | SAMEA2581294 | J2a1 | 13.7 |
| HGDP00549 | [18] | SAMEA2581681 | M1 | 14.0 |
| HGDP01009 | [18] | SAMEA2581745 | Q1a2a1a1 | 16.4 |
| HGDP00136 | [18] | SAMEA2581321 | R1a1a1b2 | 13.6 |
| HGDP00218 | [18] | SAMEA2581199 | R2a | 15.4 |
| HGDP01298 | [18] | SAMEA2581140 | L1a2 | 14.5 |
| HGDP01192 | [18] | SAMEA2581001 | N1 | 14.1 |
| HGDP01225 | [18] | SAMEA2581048 | O2a2b1a1a | 13.9 |
| HGDP01190 | [18] | SAMEA2580999 | O1b1a1a1a | 15.5 |

**Table S2:** The mutation rates required to explain the observed differences in branch length.

| Individual 1 | Individual 2 | Initial mutation rate [/bp/year] | Branch length difference [%] | Adjusted mutation rate [/bp/year] |
| --- | --- | --- | --- | --- |
| O1b1a1a1a | B-M181 | $7.34 \times 10^{-10}$ | 23.13 | $5.64 \times 10^{-10}$ |
| B-M181 | A0a1 | $5.64 \times 10^{-10}$ | 7.49 | $5.22 \times 10^{-10}$ |
| A0a1 | A00 | $5.22 \times 10^{-10}$ | 9.28 | $4.74 \times 10^{-10}$ |
| A00 | Mezmaiskaya2 | $4.74 \times 10^{-10}$ | 6.84 | $4.41 \times 10^{-10}$ |

**Table S3:** Results of mapping to the chimpanzee Y chromosome.

| Individual | Covered positions | Mean coverage | Heterozygotes | Private derived mutations |
| --- | --- | --- | --- | --- |
| Shum Laka | 3,272,603 | 7.79 | 1,301 | 653 |
| Ust'Ishim | 3,474,231 | 7.45 | 1,029 | 700 |

**Table S4:**
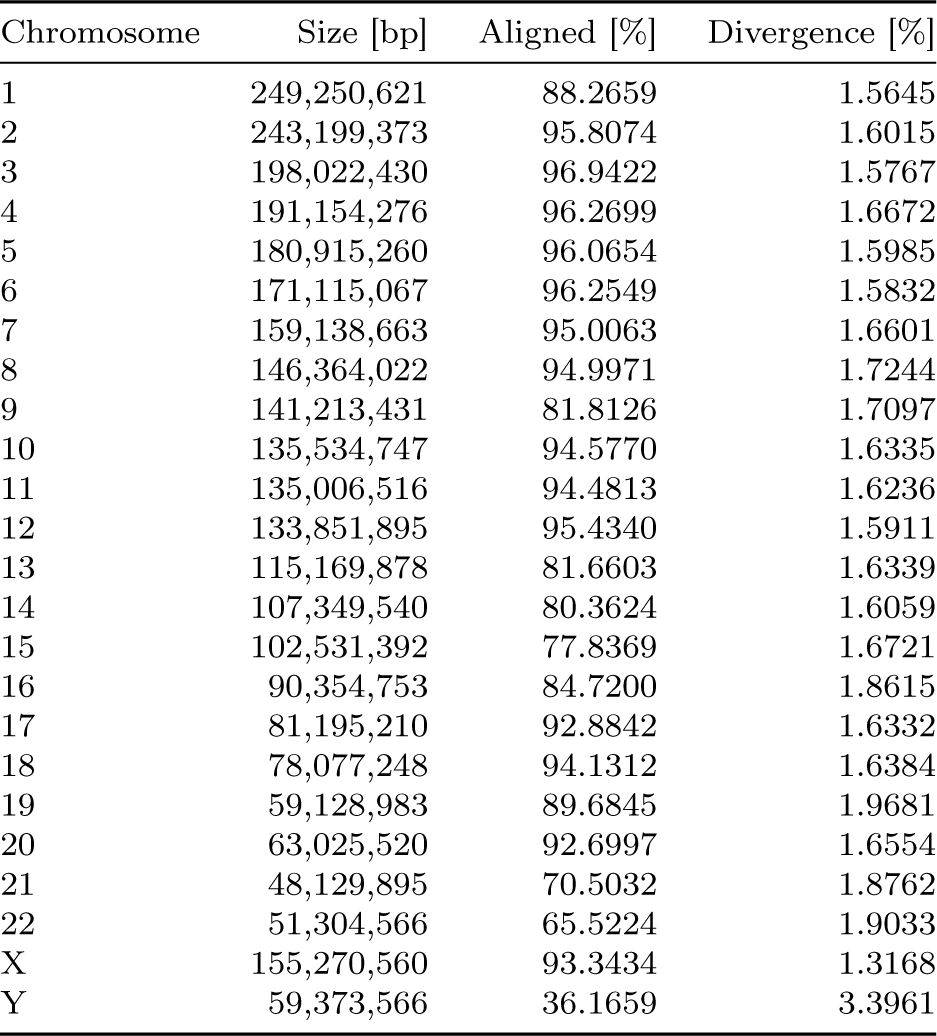
Human-chimp sequence divergence across the human genome.

| Chromosome | Size [bp] | Aligned [%] | Divergence [%] |
| --- | --- | --- | --- |
| 1 | 249,250,621 | 88.2659 | 1.5645 |
| 2 | 243,199,373 | 95.8074 | 1.6015 |
| 3 | 198,022,430 | 96.9422 | 1.5767 |
| 4 | 191,154,276 | 96.2699 | 1.6672 |
| 5 | 180,915,260 | 96.0654 | 1.5985 |
| 6 | 171,115,067 | 96.2549 | 1.5832 |
| 7 | 159,138,663 | 95.0063 | 1.6601 |
| 8 | 146,364,022 | 94.9971 | 1.7244 |
| 9 | 141,213,431 | 81.8126 | 1.7097 |
| 10 | 135,534,747 | 94.5770 | 1.6335 |
| 11 | 135,006,516 | 94.4813 | 1.6236 |
| 12 | 133,851,895 | 95.4340 | 1.5911 |
| 13 | 115,169,878 | 81.6603 | 1.6339 |
| 14 | 107,349,540 | 80.3624 | 1.6059 |
| 15 | 102,531,392 | 77.8369 | 1.6721 |
| 16 | 90,354,753 | 84.7200 | 1.8615 |
| 17 | 81,195,210 | 92.8842 | 1.6332 |
| 18 | 78,077,248 | 94.1312 | 1.6384 |
| 19 | 59,128,983 | 89.6845 | 1.9681 |
| 20 | 63,025,520 | 92.6997 | 1.6554 |
| 21 | 48,129,895 | 70.5032 | 1.8762 |
| 22 | 51,304,566 | 65.5224 | 1.9033 |
| X | 155,270,560 | 93.3434 | 1.3168 |
| Y | 59,373,566 | 36.1659 | 3.3961 |

**Table S5:**
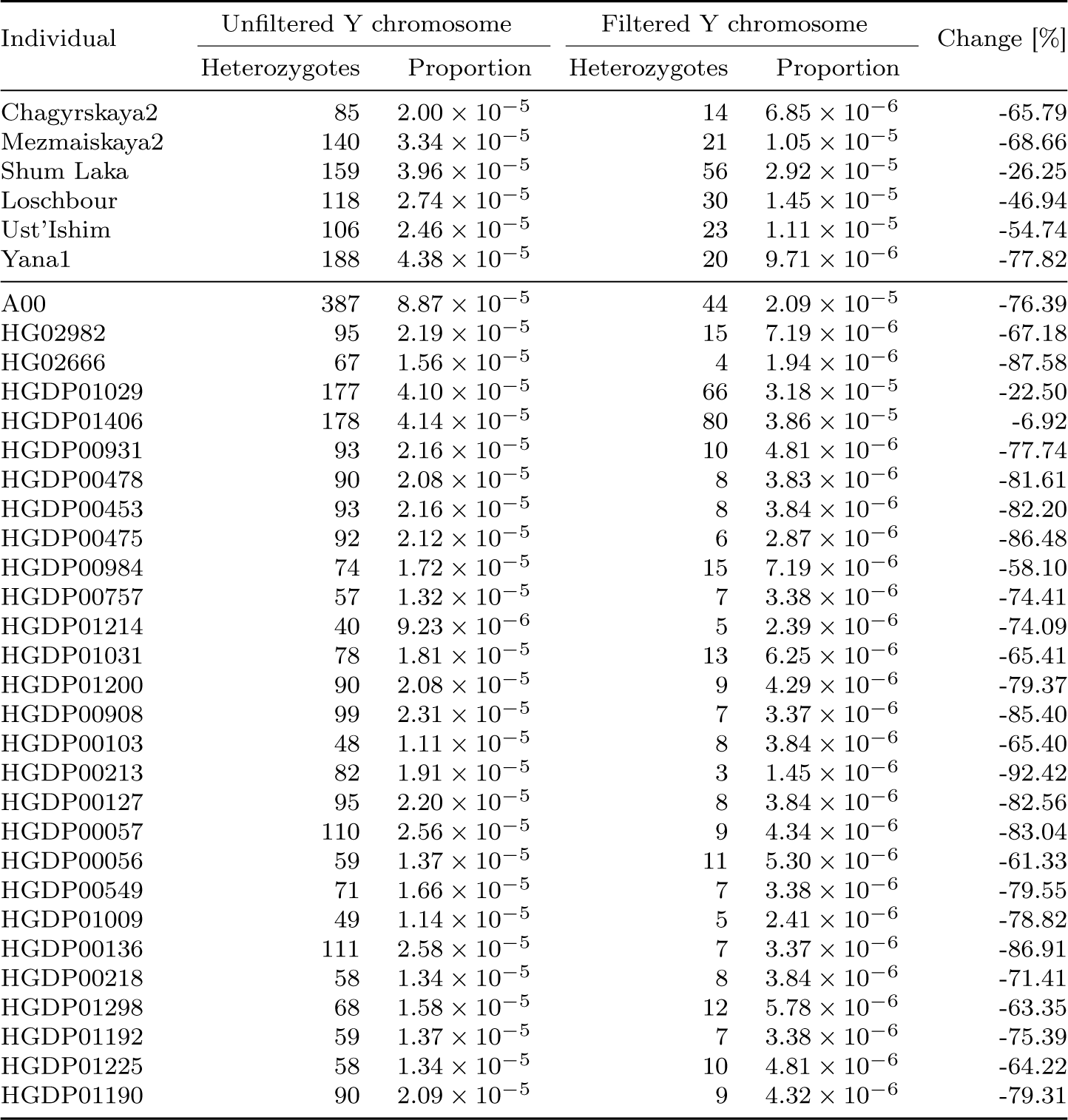
Comparing the proportion of snpAD-called heterozygotes (likely mapping errors) for the uniquely mappable part of the Y chromosome and for the uniquely mappable, divergence-filtered Y chromosome (based on a human-chimp sequence divergence cutoff of 1.9%).

**Table S6:**
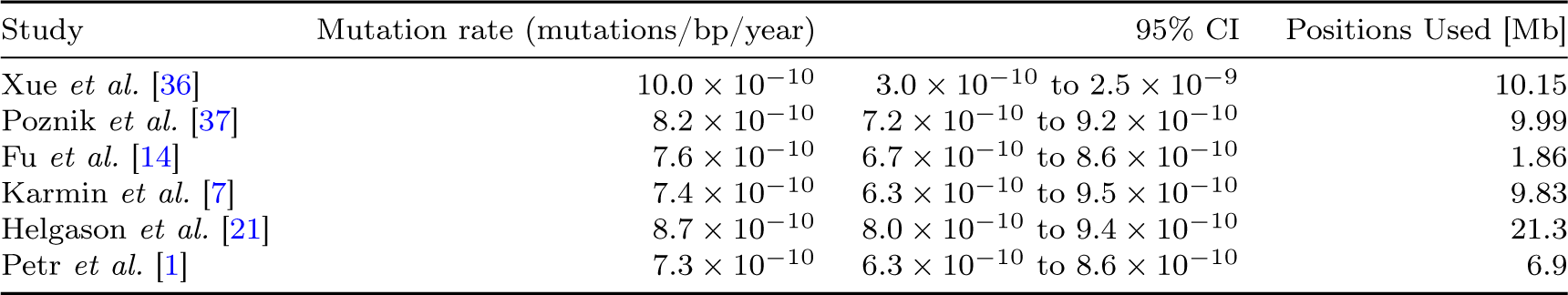
Y chromosome mutation rates estimated in previous studies.

**Table S7:** Marginal log-likelihood for the model combinations tested using the path sampling approach as implemented in BEAST2.

| Clock model | Tree model | Marginal log-likelihood |
| --- | --- | --- |
| Strict | Coalescent Constant Population | -3,203,344.16 |
| Strict | Coalescent Bayesian Skyline | -3,203,333.40 |
| Relaxed Log Normal | Coalescent Constant Population | -3,203,346.34 |
| Relaxed Log Normal | Coalescent Bayesian Skyline | -3,203,332.62 |

**Table S8:** Nucleotide substitution rates estimated by the BEAST bModelTest package.

| Substitution rate |
| --- |
| $r_{AC} = 0.548$ |
| $r_{AG} = 1.853$ |
| $r_{AT} = 0.266$ |
| $r_{CG} = 0.933$ |
| $r_{CT} = 1.853$ |
| $r_{GT} = 0.548$ |

